# How to design an optimal sensor network for the unfolded protein response

**DOI:** 10.1101/396614

**Authors:** Wylie Stroberg, Hadar Aktin, Yonatan Savir, Santiago Schnell

## Abstract

Cellular protein homeostasis requires continuous monitoring of stress in the endoplasmic reticulum (ER). Stress detection networks control protein homeostasis by mitigating the deleterious effects of protein accumulation, such as aggregation and misfolding, with precise modulation of chaperone production. Here, we develop a coarse model of the unfolded protein response in yeast, and use multi-objective optimization to determine which sensing and activation strategies optimally balance the trade-off between unfolded protein accumulation and chaperone production. By comparing a stress-sensing mechanism that responds directly to the level of unfolded protein in the ER to a mechanism that is negatively regulated by unbound chaperones, we show that chaperone-mediated sensors are more efficient than sensors that detect unfolded proteins directly. This results from the chaperone-mediated sensor having separate thresholds for activation and deactivation. Lastly, we demonstrate that a sensor responsive to both unfolded protein and unbound chaperone does not further optimize homeostatic control. Our results suggest a strategy for designing stress sensors and may explain why BiP-mitigated ER stress sensing networks have evolved.

## Background

The unfolded protein response (UPR) is a multifaceted cellular response to excess unfolded or misfolded proteins within the endoplasmic reticulum (ER) (Liu *et al.*, 2003; Schröder and Kaufman, 2005), a state referred to as ER stress. For moderate stress levels, the cellular response aims to restore protein homeostasis to the ER by upregulating quality-control enzymes and chaperones, altering ER size and shape, and attenuating translation (Harding *et al.*, 2002). When these responses fail to mitigate stress, the cell initiates apoptosis. Overloading and malfunction of the UPR are associated with numerous diseases (Kaufman, 2002; Wang and Kaufman, 2012), including diabetes (Scheuner and Kaufman, 2008; Eizirik and Cnop, 2010), cancer (Vandewynckel *et al.*, 2013), and neurodegenerative diseases (Scheper and Hoozemans, 2015; Hetz and Saxena, 2017).

A critical aspect of the UPR is the mechanism through which stress in the ER is detected and transduced to the nucleus. In the mammalian UPR, three transmembrane proteins, Ire1(inositol-requiring enzyme 1), PERK (protein kinase RNA-like ER kinase), and ATF6 (activating transcription factor 6) direct the response through three different pathways (Ron and Walter, 2007; Gardner *et al.*, 2013). Ire1 upregulates ER-localized chaperones, including the most prevalent ER protein BiP (Kar2 in yeast), and proteins involved in membrane remolding and ER-associated degradation by promoting the splicing of X-box binding protein 1 (XBP-1, Hac1 in yeast), a potent transcription factor. PERK phosphorylates the translation initiation factor eIF2*α*, which leads to an overall reduction in mRNA translation, and upregulates the transcription factor ATF4, which promotes downstream UPR genes including *Chop*, a transcription factor gene controlling apoptosis. While both Ire1 and PERK signal through similar mechanism based on the activation of kinase domains in the cytoplasmic regions, ATF6 signaling is initiated by transport of ATF6 to the Golgi, where it is processed by site-1 and site-2 proteases. The processed amino terminus then diffuses to the nucleus where it in regulates several UPR target genes, many of which overlap with those controlled by XBP-1. While mammalian cells have three interacting signaling pathways associated with the UPR, yeast possess only the Ire1 pathway (Ron and Walter, 2007; Gardner *et al.*, 2013).

As the downstream responses of the various pathways have become clearer, significant questions regarding the sensory mechanisms of Ire1, PERK, and ATF6 within the ER lumen have been raised. For Ire1 and PERK, early evidence suggested that the chaperone BiP might negatively regulate the activation of the sensory molecules (Kimata *et al.*, 2003). However, a mechanism involving only BiP was shown to be insufficient since the UPR is inducible in cells with modified Ire1 and PERK that are incapable of binding BiP (Kimata *et al.*, 2004). This led to the hypothesis that the activation of sensory proteins by unfolded protein ligands is buffered by BiP (Kimata *et al.*, 2004, 2007; Pincus *et al.*, 2010). Further support for this hypothesis comes from structural similarities between Ire1 luminal domain dimers and major histocompatibility complexes, which both show a favorable groove for direct peptide binding (Credle *et al.*, 2005). Additional studies have provided evidence of unfolded protein interaction with Ire1 and PERK in yeast (Gardner and Walter, 2011), and more recently with human Ire1 (Karagöz *et al.*, 2017). Other evidence suggests that BiP binding to Ire1 and PERK may be allosterically regulated by unfolded proteins (Carrara *et al.*, 2015), providing an alternative mechanism of activation. Since evidence suggests that Ire1 and PERK bind an overlapping, but distinct set of proteins as BiP (Karagöz *et al.*, 2017), it seems possible that both of these mechanism are realized in vivo.

While the molecular details of the activation mechanism have yet to be fully resolved, one fact is clear: stress-sensing is quite complex. Our aim in this work is to better understand why such a complex system might be beneficial for stress detection, and thereby provide insight into the evolutionary forces guiding stress-sensor design. Specifically, we ask: How do phenotypic features of a sensory network affect the trade-off between accumulation of unfolded proteins and metabolically-efficient chaperone production in response to stress? We start from the naïve perspective that the simplest way to detect stress would be to directly count the number of unfolded proteins within the ER. Using a coarse-grained model of the UPR based on direct sensing of unfolded proteins, we determine the optimal shape of response functions for acute and chronic stress conditions. Next, following experimental evidence that chaperones inhibit stress-sensor activation, we determine the optimal response function for acute and chronic stress when sensor activation is negatively regulated by freely available chaperone. Comparing the optimal performance of sensory systems that directly measure unfolded protein concentration with those that respond instead to available chaperone, we show that indirectly measuring stress through free chaperone concentration leads to a more efficient use of chaperone in mitigating stress. Finally, we consider whether further benefits can be obtained by combining both sensing modalities, as is observed experimentally.

## Methods

### Model Description

To model the UPR (Figure 1), we develop a course model in which we consider only two species explicitly: free unfolded client proteins, *U*, and the total number of a generic chaperone present in the ER, *C*_*T*_. Descriptions of the UPR at a similar level of detail have been previously used to investigate the benefits of translational regulation (Axelsen and Sneppen, 2004; Trusina *et al.*, 2008). *C*_*T*_ encompasses both free chaperone, *C*_*F*_, and chaperone forming a folding complex with client proteins, *C · U*, such that *C*_*T*_ *= C*_*F*_ + *C · U*. The unfolded proteins represent those proteins in the ER with a significant number of exposed hydrophobic residues, and hence are highly active and aggregation prone. The chaperones mediate this activity by restoring the proteins to the folding pathway. Proteins returned to this pathway are assumed to fold without incident and are secreted from the ER. The governing delay differential equations for the reactive unfolded protein and total chaperone levels are

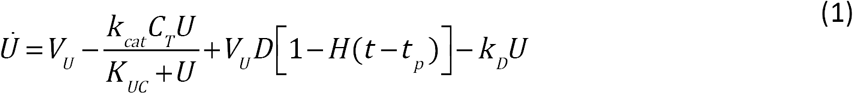

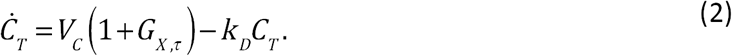

**Figure 1:**
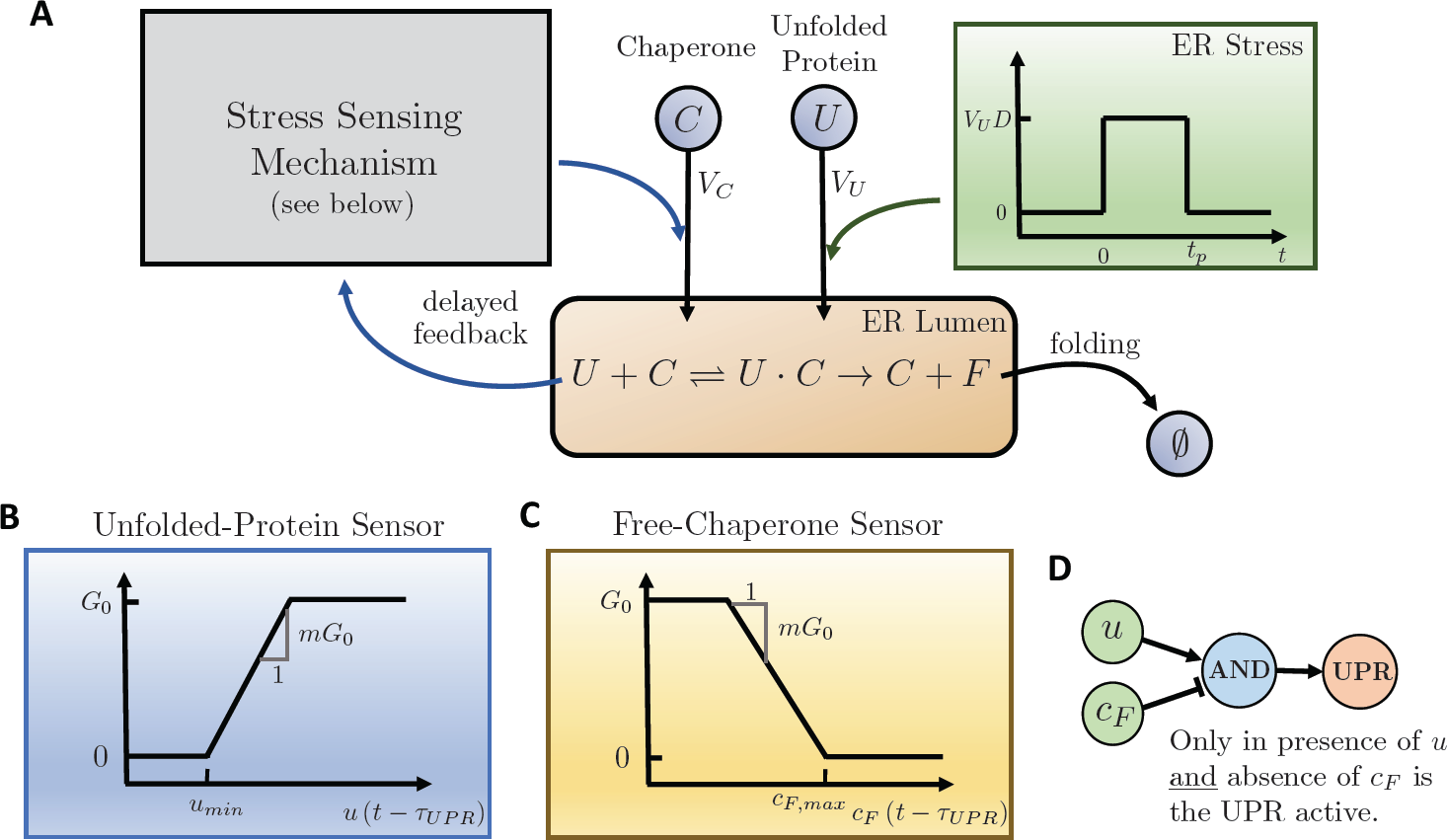
UPR model schematic. Panel A shows a schematic overview of the UPR model. Protein production in the ER lumen is modeled as a chaperone-assisted folding process that returns unfolded proteins to the folding pathway (*F*), which are removed from the system. The basal flux of unfolded proteins into the ER, *V*_*U*_, is augmented in the case of stress by an additional *V*_*U*_*D* flux. The pulsatile stress (top, right box) lasts for duration *t*_*p*_. Upon increased accumulation of unfolded protein in the ER, the UPR (top, left box) is activated, upregulating the influx of chaperone into the ER to mitigate the increased folding load. Specific stress-sensing models are shown in panels B-D. Panel B depicts a UPR activation model that responds directly to the concentration of unfolded protein in the ER lumen in a piecewise-linear manner with slope proportional to *m*, and activation threshold *u*_*min*_. Panel C shows a chaperone-mediated sensor model that is activated when the concentration of free chaperone decreases below a threshold value *C*_*F,max*_. Panel D shows the logic circuit for the AND-switch, which combines the unfolded-protein sensor and the free-chaperone sensor.

In Equation (1), the first term *V*_*U*_ represents the basal flux of unfolded proteins into the ER. The second term (*k*_*c*at_*C*_*T*_*U*)*/*(*k*_*UC*_ + *U*) describes the catalytic activity of the chaperone on the protein. The third term *V*_*u*_*D*[1 - *H*(*t - t*_*p*_)] represents a state of stress, which we model as a square pulse of increased flux starting at *t =* 0, where the pulse height is *D* times the basal flux, *H* is the Heaviside function, and *t*_*p*_ is the pulse duration. The final term *k*_*D*_*U* captures the decrease in unfolded proteins present in the ER due to dilution and degradation, with *k*_*D*_ being the inverse half-life of a protein in the ER. Similarly, in Equation (2), *V*_*C*_ is the basal influx of chaperone, and the dilution and degradation of chaperone is described by the final term. *G*_*X*,*τ*_ represents the response function for the feedback response of the UPR. The subscript *X* denotes the specific model for UPR activation as described below, and the subscript *τ* indicates that the response depends on the state of the system at a previous time *t - t*_*UPR*_.

A detailed description of the model derivation and assumptions is available in the Supplemental Material. The parameterization of the model is constrained by experimental results from literature and is presented in Table 1.

**Table 1.**
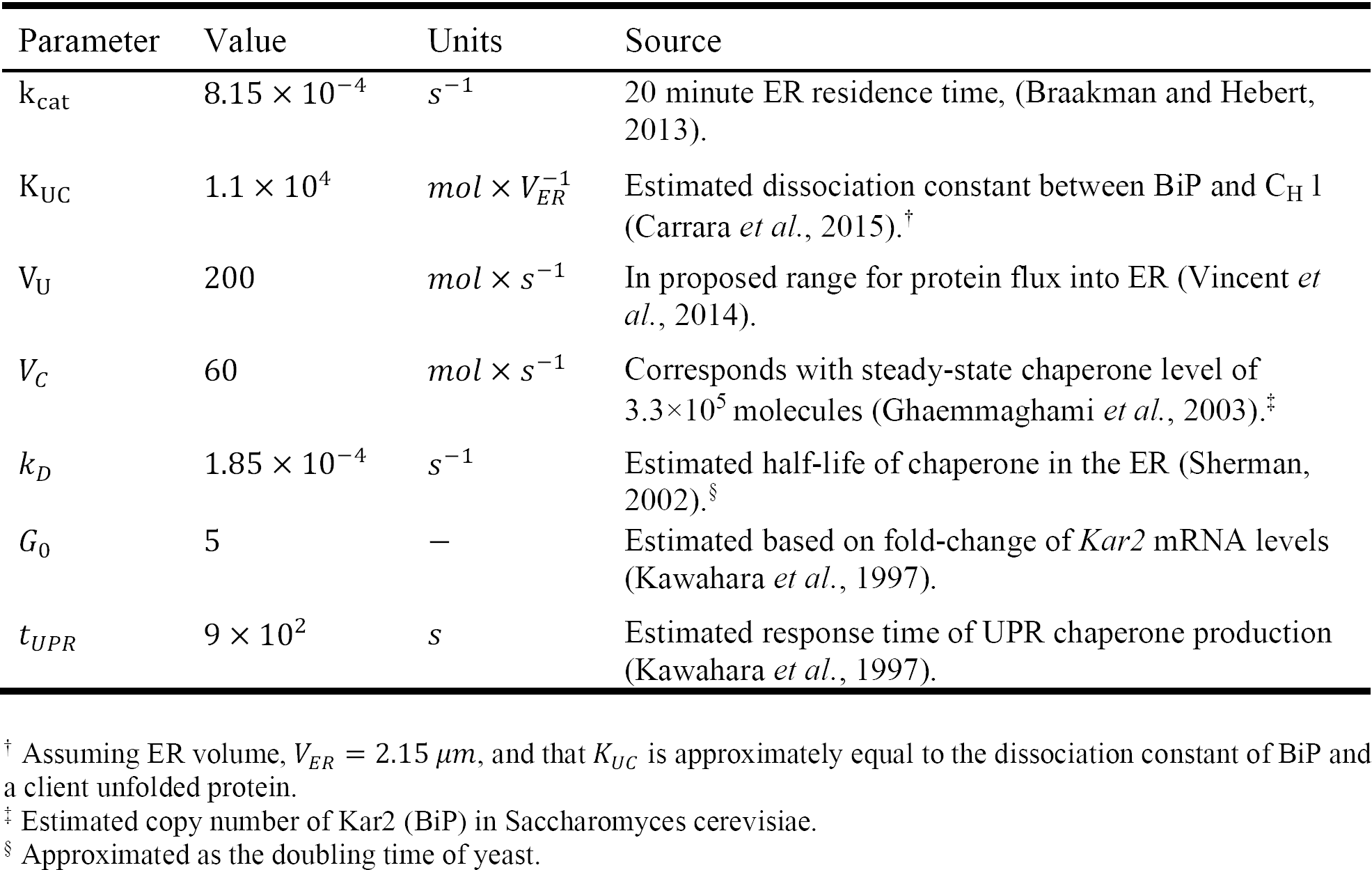
Model Parameters

| Parameter | Value | Units | Source |
| --- | --- | --- | --- |
| $k_{cat}$ | $8.15 \times 10^{-4}$ | $s^{-1}$ | 20 minute ER residence time, (Braakman and Hebert, 2013). |
| $K_{UC}$ | $1.1 \times 10^4$ | $mol \times V_{ER}^{-1}$ | Estimated dissociation constant between BiP and C<sub>H</sub> 1 (Carrara *et al.*, 2015).<sup>†</sup> |
| $V_U$ | 200 | $mol \times s^{-1}$ | In proposed range for protein flux into ER (Vincent *et al.*, 2014). |
| $V_C$ | 60 | $mol \times s^{-1}$ | Corresponds with steady-state chaperone level of $3.3 \times 10^5$ molecules (Ghaemmighami *et al.*, 2003).<sup>‡</sup> |
| $k_D$ | $1.85 \times 10^{-4}$ | $s^{-1}$ | Estimated half-life of chaperone in the ER (Sherman, 2002).<sup>§</sup> |
| $G_0$ | 5 | — | Estimated based on fold-change of *Kar2* mRNA levels (Kawahara *et al.*, 1997). |
| $t_{UPR}$ | $9 \times 10^2$ | $s$ | Estimated response time of UPR chaperone production (Kawahara *et al.*, 1997). |

<sup>†</sup> Assuming ER volume, $V_{ER} = 2.15 \mu m$ , and that $K_{UC}$ is approximately equal to the dissociation constant of BiP and a client unfolded protein.
<sup>‡</sup> Estimated copy number of Kar2 (BiP) in *Saccharomyces cerevisiae*.
<sup>§</sup> Approximated as the doubling time of yeast.

#### Non-dimensionalization of model equations

To facilitate analysis, we non-dimensionalize the model by choosing the degradation time 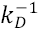 to be a characteristic timescale and *V*_*c*_ /*k*_*D*_ to be a characteristic concentration. This leads to a normalized form of the model

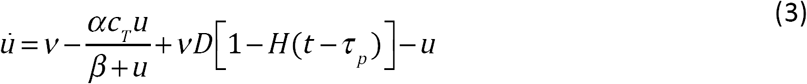

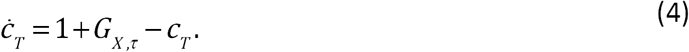

In addition to the normalized times for the pulse duration *τ*_*p*_ *= t*_*p*_*k*_*D*_ and UPR response time *τ*_*UPR*_*= t*_*UPR*_*k*_*D*_, we have introduced three dimensionless parameters

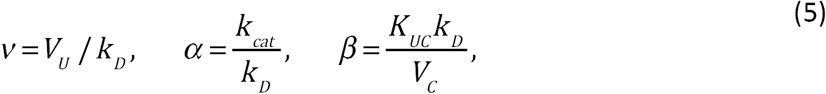

Where *v* is the dimensionless influx rate of unfolded protein into the ER, α is the dimensionless catalytic constant for chaperone-assisted folding, and β is the dimensionless Michaelis constant for the chaperone-unfolded protein interaction.

### Stress sensing mechanisms

In yeast, the level of stress in the ER is sensed by the transmembrane protein Ire1, which then facilitates the splicing of the transcription factor Hac1. Hac1 upregulates the transcription of *Kar2*, among other genes, increasing the translation of the ER chaperone Kar2 (BiP). We capture this process through a phenomenological model governing the level of activation of Ire1. Since there is a finite amount of time required for splicing, transcription, and translation, the increased flux of chaperone depends on the level of activity of Ire1 a time *τ*_*UPR*_ before. We incorporate this lag time as a time delay on the state variables *u* and *C*_*T*_ in the response function, and denote 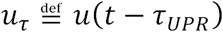 and 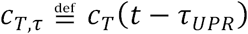. We consider three different response functions, each representing a different stress-sensing mechanism.

#### Unfolded-protein sensor

In the first mechanism, activation is directly related to the concentration of unfolded protein in the ER (Figure 1B). In this case, the activation function, *G*_*u,τ*_ is

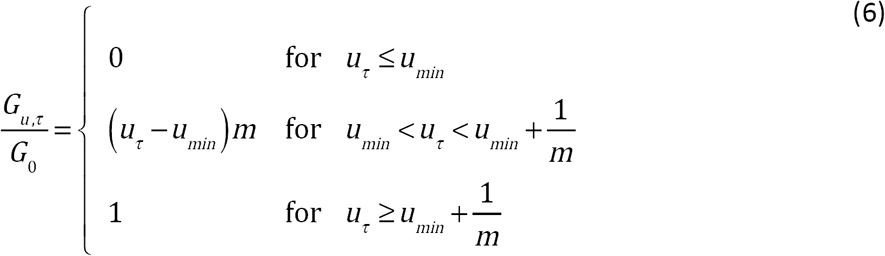

Where *G*_0_ the maximal gain, *u*_*min*_ is the level of unfolded protein for which the UPR first activates, and *m* is the slope of response function. The inverse of *m* can be interpreted as the width of the active range of the sensor. The response function resembles common sigmoidal responses found in biology, but has the advantage of having the same steady state for any parameterization of the response function so long as the steady state is below the activation threshold. We refer to this mechanism as the “*U*-switch" mechanism.

#### Chaperone sensor

The second mechanism we consider is one in which activation of the UPR is inhibited by free chaperones in the ER (Figure 1C). In this case, the activation function, *G*_*c,τ*_, is

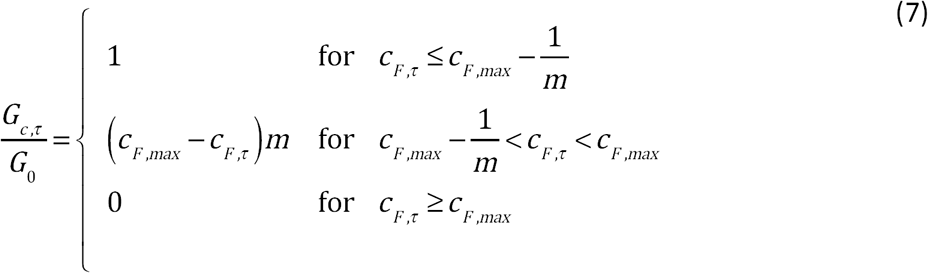

where *C*_*F,τ*_ is the concentration of free chaperone at time *t - τ*_*UPR*_, which is given by

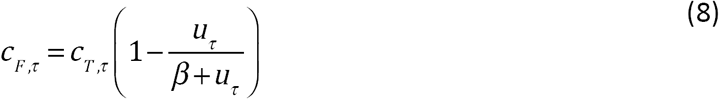

(see Supplemental Material for a detailed derivation). *C*_*F,max*_ is the threshold concentration of free chaperone above which there is no activation, and *m* is the slope of the response. We refer to this mechanism as the “*C*_*F*_-switch" mechanism.

#### Unfolded-protein-AND-chaperone sensor

The final mechanism consists of a combination of the previous two mechanisms such that both the unfolded-protein-sensing function and the chaperone-sensing function must be active, that is *u* concentration must be high and *C*_*F*_ concentrations low, for the UPR to respond (Figure 1D). The activation function for this case, *G*_*AND,τ*_, is

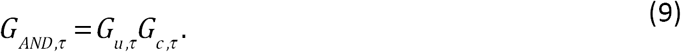

We refer to this mechanism as the “AND-switch" mechanism as it becomes a logical AND function in the case that the unfolded-protein and chaperone sensors become discrete on-off switches.

## Computational Approach

### Fitness measures and Pareto optimization

It is often the case that the fitness of a phenotype is a function of several independent quantities. While the UPR influences cellular fitness in a wide-ranging set of interactions and functions that remain to be fully understood, we choose to focus on two measures of fitness that are central to UPR function: (i) the amount of excess unfolded protein present in the ER while the cell is under stress, and (ii) the total excess production of chaperone in response to an impulse of stress. The first measures how effectively the UPR is able to mitigate the negative effects of high concentrations of highly-reactive protein species (such as aggregation and misfolding) in the ER. The second measures the metabolic cost associated with rapidly reducing stress through chaperone production. When dealing with multiple fitness functions, one option is to choose a weighting for each and use the weighted sum of the individual fitness functions as a scalar measure of overall fitness. However, this often requires an ad hoc choice of weights, making the results somewhat subjective. An alternative approach is to use Pareto optimization, which seeks to determine Pareto-efficient solutions in fitness space. The set of Pareto-efficient solutions, also called the Pareto front, is the set of points in fitness space for which any improvement in one fitness measure can only be achieved by a decline in another fitness measure. Any specific weighting scheme in a scalar weighted-sum fitness measure corresponds to a point on the Pareto front. This technique has been used to probe phenotype space distributions in general (Savir *et al.*, 2010; Shoval *et al.*, 2012), as well as being applied to several biological problems in particular, including gene regulatory networks (Warmflash *et al.*, 2012) and homeostatic control (Szekely *et al.*, 2013).

The fitness landscape guiding the evolution of the cellular stress response very likely has many competing factors, such as the response time or noise reduction, in addition to the accumulation of unfolded proteins in the ER lumen and the production of mitigating chaperones. Here, we focus only on the trade-off between unfolded protein concentration and chaperone production as these are two fundamental features of homeostatic UPR control. We note that this trade-off is only one of many possible guiding principles in stress-sensor design. The function measuring the cost of excess unfolded protein accumulation in the ER is given by

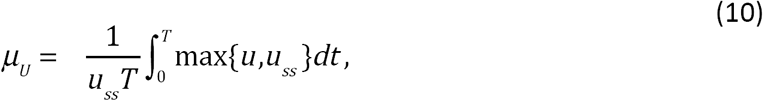

where *T* is the duration of the simulation, max{*u, u*_*ss*_} is the maximum between the unfolded protein level and the steady-state unfolded protein concentration at each time, and the prefactor to the integral is a normalization constant. Note that we seek to minimize a cost function. The corresponding fitness function would be the negative of the cost function and is maximized. μ_*U*_ measures the time-average of the excess unfolded protein in the ER, normalized by the steady-state unfolded protein concentration. The cost function for chaperone production is computed by

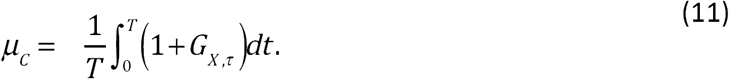

*μ*_*C*_ gives the time-average of the chaperone production rate. Together, the measures (*μ*_*U*_, *μ*_*C*_) form the two-dimensional fitness space for the UPR.

In the case of the *U*-switch and *C*_*F*_-switch, the Pareto fronts are computed using a brute-force method by calculating the objective functions over a 100-by-100 grid of logarithmically-spaced points in the ranges *m* ∈ [0.1,1000], *u*_*min*_ ∈ [0.01*u*_*ss*_,1000*u*_*ss*_], and *C*_*F,max*_ ∈ [0.001*c*_*F,ss,*_,0.999*C*_*F,ss*_]. We found that using a brute-force method provides efficient coverage of the extreme points on the Pareto front. For the AND-switch, in which the number of independent parameters is four as opposed to two, the Pareto front is calculated using the Non-dominated Sorting Algorithm II (Deb *et al.*, 2002) as implemented in the Python software platypus (Hadka, 2015).

### Numerical solution of delay differential equations

To simulate Equations (3) and (4), we use the Python software pydelay, which implements the Bogacki-Shampine method to compute trajectories of systems of delay differential equations (Flunkert and Schoell, 2009). Solutions to delay differential equations require the specification of a time history for each variable for at least as long as the longest delay present in the system. Here, we use the steady-state of the unfolded protein and chaperone levels for the time history such that our perturbations due to stress are deviations from the steady functioning of the ER folding machinery. The steady states are found by setting Equations (3) and (4) equal to zero under the assumption that the UPR is operating at a basal level below the threshold for activation, which leads to

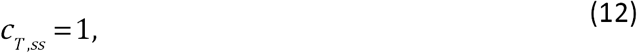

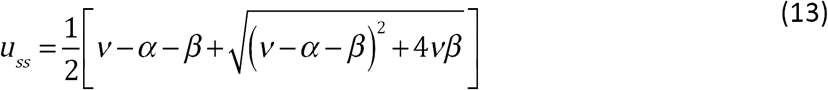

These equations are valid only so long as they are consistent with the assumption that the UPR is inactive at the steady state, i.e. *u*_*ss*_ < *u*_*min*_ for the *U*-switch and *C*_*F,ss*_ *> C*_*F,max*_ for the *C*_*F*_- switch and both conditions for the AND-switch. This requirement places constraints on the values taken by *u*_*min*_ and *C*_*F,max*_ for a given parameterization. For the optimization procedure described below, we ensure that the steady state for the baseline influx does not activate the UPR for all simulations. Additionally, if the slope of the activation function, *m*, for the *C*_*F*_-switch is small enough, the activation function can be less the *G*_0_ when *C*_*F*_ *=* 0, effectively lowering the maximum possible response. To avoid this, we also ensure that both the *C*_*F*_- and AND-switches can reach maximum activation for positive *C*_*F*_. If the amplitude of the stress is small enough, the existing (steady-state) chaperone concentration is sufficient to maintain a new steady-state concentration of unfolded protein that is still below the activation threshold for the UPR (see Supplemental Material). Conversely, for a fixed stress amplitude, *D*, a maximal value of the activation threshold can be determined, beyond which no response will occur. This provides a boundary constraint for the optimization procedure outlined in the previous section. Despite starting the simulations with a history at the steady state, numerical artifacts can lead occasionally to small fluctuations at the onset of the simulations. To ensure these do not interact with the prescribed perturbations we seek to analyze, each simulation is allowed to relax for a period of 3*pτ*_*UPR*_before the pulse of stress is applied. Following the equilibration period, each model is simulated for time of *τ*_*p*_ + 30*τ*_*UPR*_. This allows all simulations, regardless of pulse shape and response function parameterization to return to a steady state following the stress response.

### Code Availability

Python scripts used to simulate the model and calculate Pareto sets can be found at https://bitbucket.org/schnell-lab/upr_feedback_control/src/master/. All codes were run using Python version 2.7.

## Results

Any effective stress sensing network must be sensitive to the concentration of unfolded proteins in the ER lumen. However, the efficiency with which a sensor controls the UPR will depend on the time course of the stress. With this in mind, we consider the response of sensory networks to two types of characteristic stress time courses: (i) a sustained chronic stress in which the system adjusts to a new steady-state at a larger-than-usual protein influx, (ii) acute stresses of varying amplitude and duration, in which the on- and off-dynamics of the UPR become important. For each stress type, we determine the set of Pareto optimal sensor designs for the unfolded-protein sensor and the free-chaperone sensor, and compare their features and efficiency. Lastly, we compare the *U*-switch and *C*_*F*_-switch sensor models individually with the combined AND-switch model. Results from the response of these three sensing strategies to different stresses provides insights into the potential evolutionary benefits of sensor network design.

### Optimal design of a sensor for chronic stress

We initially consider a cell subjected to a sustained pulse of increased unfolded protein translocation rate into the ER. The stimulus, which we call chronic stress, is a step increase of influx rate that continues indefinitely, allowing the system to fully acclimate to the stressed state. For this stress signal, we have computed the Pareto-optimal parameterizations of both the unfolded-protein-sensing and the free-chaperone-sensing mechanisms (Figure 2A). In the case of chronic stress, both objective functions are determined by the new steady-state reached by the UPR-activated system. Both mechanisms can reach the same set of steady-states, and hence have the same Pareto fronts. Additionally, for each mechanism, the switching function that produces a steady state is not unique. In fact many combinations of slope and threshold will lead to the same steady-state in response to chronic stress, as shown schematically in Figure 2A (insets). This can be readily seen by considering two switch parameterization (*u*_*min,*1_,*m*_1_) and (*u*_*min,*2_,α*m*_1_) in which the slope of the second response function is related to the slope *m*_1_ through a linear scaling α. In the linear regime of the response, the steady-state chaperone and unfolded protein levels, 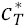 and *u*\* are related through the equations

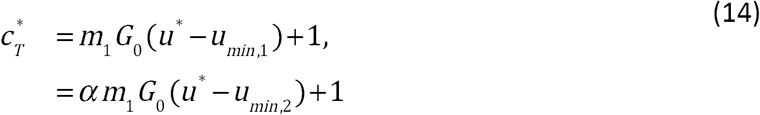

which leads to the linear relationship between *u*_*min,*1_ and *u*_*min,*2_ in terms of α

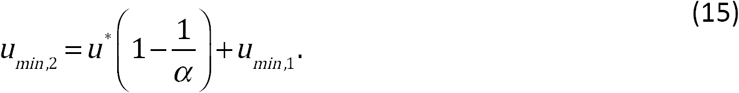

**Figure 2:**
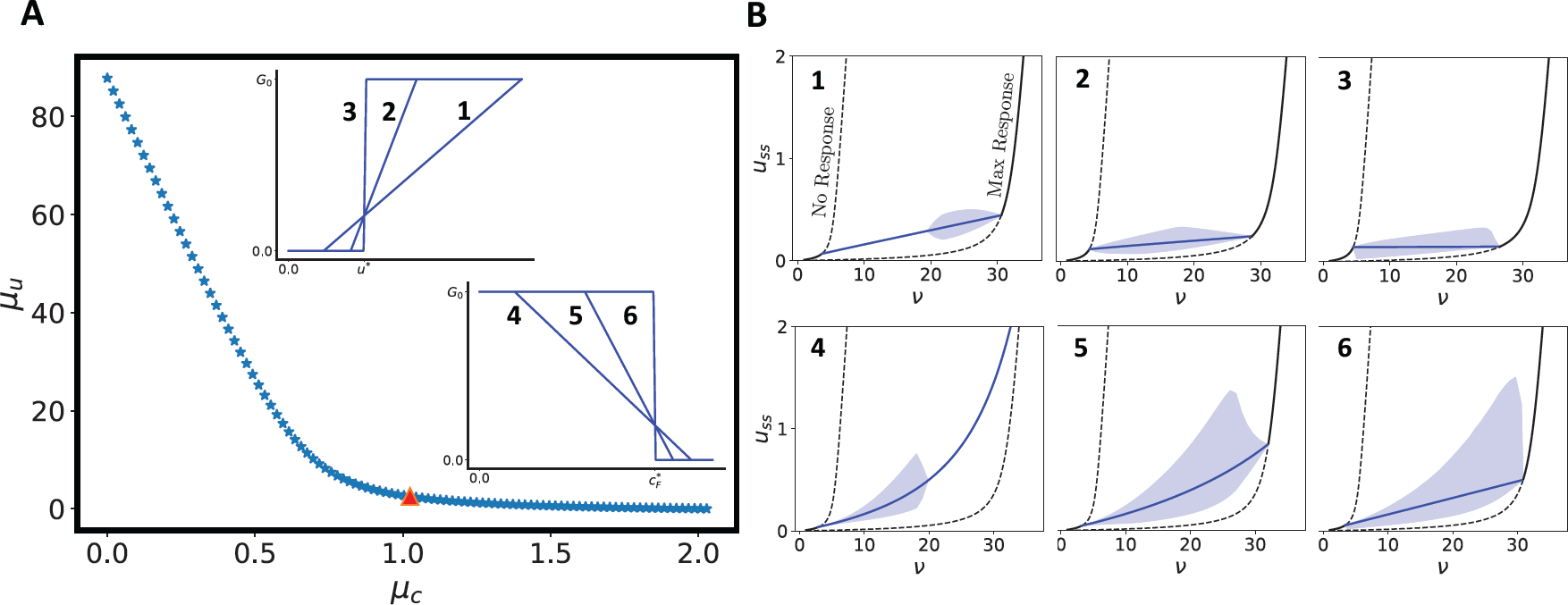
Pareto front for chronic stress response. Panel A shows the Pareto front (blue stars) for a sustained stress, where the system adapts to an increased stress level. The inset provides representative activation functions for the point on the Pareto front marked by a red triangle for the *U*-switch (top inset) and *C*_*F*_-switch (bottom inset). Panel B shows the steady-state unfolded protein level as a function of influx rates. The black lines show the steady-states of the system when there is no UPR activation (left) and full UPR activation (right). The solid and dashed sections of the lines correspond to stable and unstable fixed points, respectively. The blue lines show the analytically-determined fixed points for intermediate activation of the UPR. The shaded regions show the range of oscillations of numerically-simulated solutions. The numbers in the top center of each panel correspond to the activation functions shown as insets in Panel A.

Hence, infinitely many combinations of (*u*_*min*_, *m*) will lead to the same steady state and the same efficiency in dealing with a specific state of chronic stress. The same is true for the chaperone-based sensor.

However, the sensitivity of the response to changes in stress and the damping of oscillations depend on the particular parameterization of the switching function. As the response becomes steeper, Figure 2B shows that the mean level of unfolded protein changes less in response to the increases in influx. However, this comes at the cost of oscillations. Thus, while tighter control over the mean value of protein concentration can be achieved with a more abrupt response, a more graded response allows the system to adjust to a range of different levels of chronic stress without oscillations.

### Efficient sensor design for acute stress

In the case of chronic stress, the quality of a response depended on the steady-state behavior of the stressed system. The dynamics of system were not important, except in regard to oscillations in systems with steep responses. In contrast, for shorter stress events in which a new steady state may not be reached, the dynamics of the UPR are essential in determining the efficiency of the response. To investigate this, we consider the response of the system to stress pulses of different shape. Due to the nonlinear coupling and delays, analytical solutions are either unavailable or uninsightful for the transient pulse response. Hence, we numerically determine the Pareto fronts for each mechanism across a range of pulse shapes. Figure 3A-E show Pareto fronts for a set of pulses in which the total protein influx is conserved (i.e. *Dτ*_*p*_ *= Constant*), and the amplitude and duration of the pulse is modulated. For small-amplitude, long-duration pulses the unfolded protein-based switch is slightly more efficient when excess unfolded protein accumulation is the primary cost. However, the magnitude of the difference between the two mechanisms is small relative to the difference seen for other pulses. Similarly, for very short pulses of larger amplitude, the two mechanisms are essentially equivalent. However, for intermediate pulse shapes, the chaperone-based switch can be substantially more efficient. To quantify this, we divide the area under the Pareto front of the *C*_*F*_-switch by the area under the *U*-switch Pareto front

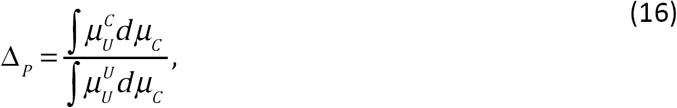

where 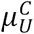 is μ_*U*_ of the unfolded protein-based mechanism and 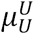 is μ_*U*_ of the chaperone-based mechanism and the integrals are computed over the range of the μ, on the Pareto fronts. When *Δ*_*P*_ *>* 1, the chaperone-based mechanism provides a more efficient response across parameterizations, while the unfolded protein-based mechanism is more efficient when *Δ*_*P*_ *<* 1. We note that the ability of this metric to quantify the degree to which one mechanism is superior to another depends on the Pareto fronts not crossing. Observation of the calculated Pareto fronts indicates that in the small range where the Pareto fronts do cross, the *C*_*F*_-switch is superior when the more dominant cost is the mitigation of stress (see Figure 3A-B). Figure 3F shows *Δ*_*P*_ for the cases *Dτ*_*p*_ *=* 1. There is a large range of intermediate pulses for which the chaperone-based switch is significantly more efficient for any choice of parameterization.

**Figure 3:**
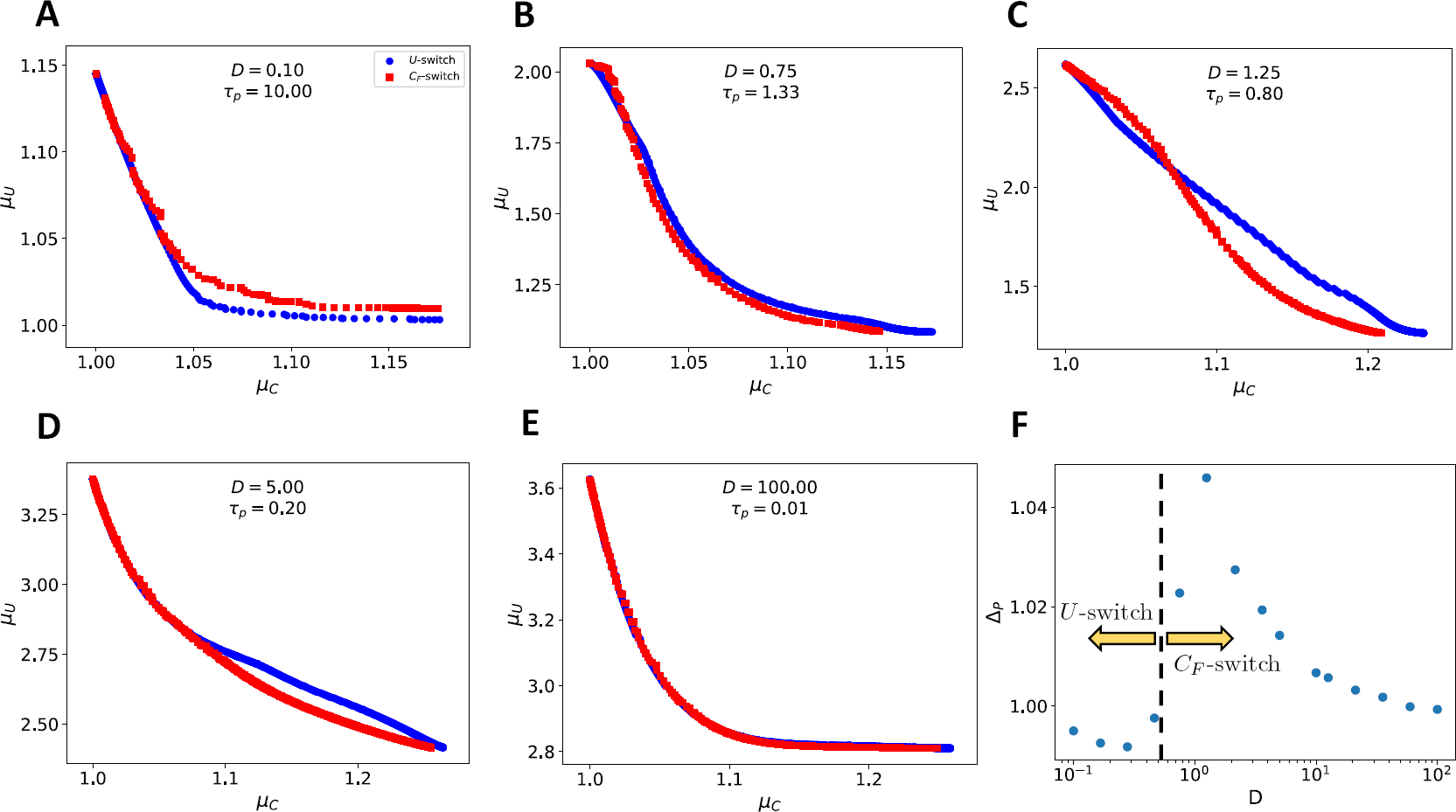
Pareto front for acute pulse. Panels A-E show the Pareto fronts corresponding to the *U*-switch sensor (blue circles) and the *C*_*F*_-switch sensor (red squares) for different pulse shapes. The total excess protein in each pulse is 1, with the pulse height and duration changing. Panel F shows the normalized area between the Pareto fronts of each mechanism as a function of pulse amplitude (with total pulse influx conserved). The black dashed line delineates the pulse shapes for which the *U*-switch is superior (left) from those for which the *C*_*F*_-switch is superior (right).

The added efficiency of the *C*_*F*_-switch can be understood by examining the phase-plane trajectories of each mechanism (Figure 4B). The projection of the unfolded-protein sensor threshold into the *C*_*T*_ - *u* plane is a horizontal line, while the free-chaperone sensor threshold projects onto a line with a positive slope (see Supplemental Material for derivation). Whereas the horizontal threshold of the unfolded-protein sensor threshold means that the UPR will turn on and off at identical levels of stress, the slope of the free-chaperone sensor means that the on- and off-thresholds are no longer symmetric with regard to stress. This imparts three advantages to the free chaperone sensing system. First, it allows for earlier activation when a stress arises, thereby reducing the maximum concentration of unfolded protein that occurs during the stress event (shown as a heat map on the Pareto fronts in Figure 4A.). Additionally, the gradient of the slope with respect to unfolded protein concentration is maximal at the onset of stress for the *C*_*F*_- switch, allowing an initially-strong response (see heat map in Figure 4C). Lastly, the system is able to deactivate the response sooner when the level of stress begins to subside, thereby preventing the excess production of unneeded chaperone, as seen in the corresponding time courses shown in Figure 4C.

**Figure 4:**
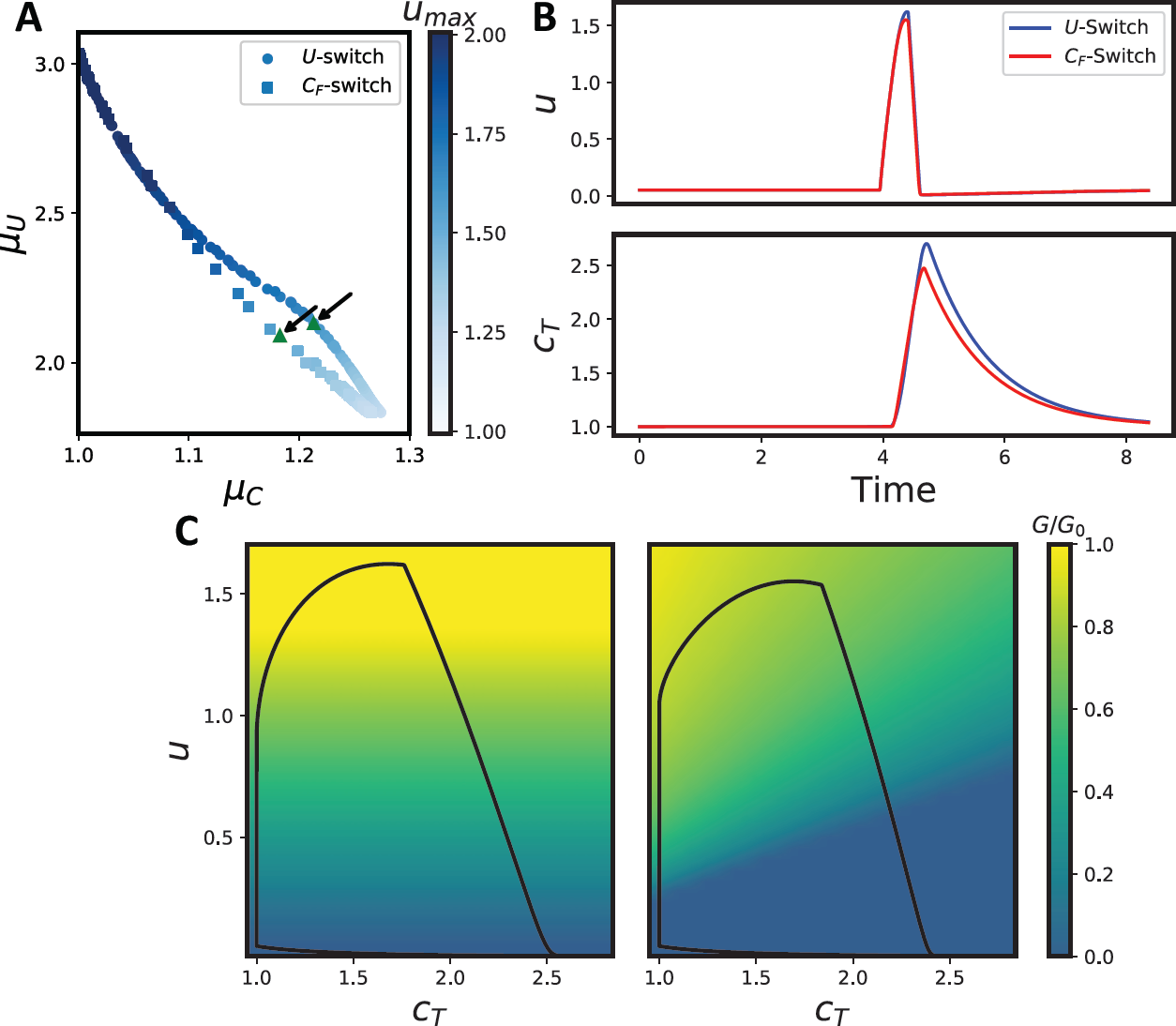
Comparison of Pareto fronts for *U*-switch and *C*_*F*_-switch. Panel A shows Pareto fronts for the UPR activated by the level of unfolded protein directly (circles) and by the level of free chaperone (squares). The chaperone-sensing system performs more efficiently than the unfolded-protein-sensing system in all cases. The coloring of the markers corresponds to the maximal unfolded protein concentration reached during the stress event. Panel B shows the time courses for unfolded protein (top) and total chaperone (bottom) during the stress event for the two Pareto-optimal models indicated by the arrows (and green triangles) in Panel A. Panel C shows the phase-plane trajectories (black curves) for the same two parameterizations as in Panel B. The background coloring indicates the activation level for the UPR in each model. Due to the slanted activation threshold of the *C*_*F*_-switch, the UPR deactivates at a higher level than the *U*-switch. Parameter values for the stress pulses are: *D =*2.15, *τ*_*p*_ *=*0.46.

While the cost function μ_*U*_ measures the integrated excess protein accumulation with the ER over the time course of the stress, the maximal level of unfolded protein may also be an important physiological measure of fitness. In Figure 4A, the Pareto fronts for each mechanism are colored corresponding to the peak unfolded protein concentrations experienced during the stress pulse. For the same amount of excess chaperone production, the chaperone-based sensor leads to both less integrated excess unfolded protein and lower peak unfolded protein for nearly the entire Pareto front, except for where the two mechanism provide equivalent responses.

### Logical AND-switch sensor combining chaperone and unfolded protein concentrations

Experimental evidence supports a model for UPR activation that relies on both the dissociation of BiP from the sensory protein, and the binding of an unfolded protein to sensor oligomers (Oikawa *et al.*, 2007; Pincus *et al.*, 2010; Karagöz *et al.*, 2017). To investigate how a sensor integrating both the concentration of unfolded protein and the concentration of free chaperone (which serves as an indirect measure of chaperone-sensor binding), we combine the models for the *U*-switch with the *C*_*F*_-switch by multiplying the two activation functions (see Equation (9)) to form the AND-switch (shown in Figure 1D). The AND-switch is zero everywhere that either the *U*- switch or the *C*_*F*_-switch are zero and fully activated only when both individual switches are also fully activated. In this way, as the two slope parameters, *m*_*u*_ and *m*_*c*_, become large, the AND-switch approximates a logical AND gate for the two input signals.

Figure 5 shows a comparison between the Pareto sets for the AND-switch and the two individual switching functions for a pulse with amplitude *D =* 1.0 and duration *τ*_*p*_ *=* 0.5. At low values of μ_*c*_ on the Pareto fonts, the AND-switch coincides with the *U*-switch, while at higher values of μ_*c*_ on the Pareto front, the AND-switch and the *C*_*F*_-switch coincide. For all cases, the AND-switch aligns with the better of the *U*-switch and the *C*_*F*_-switch, but never outperforms them. This suggests that when the trade-off between chaperone production and unfolded protein buildup favors greater chaperone production, the optimal sensing strategy is to monitor the free chaperone, as demonstrated by the *C*_*F*_-switch results. In this case, the unfolded protein concentration does not directly control UPR activation, although it may still influence the system in ways not considered here, such as stabilizing the signal or reducing noise. When the trade-off places a premium on chaperone efficiency (low μ_*C*_), the Pareto front more closely follows that of the *U*-switch than the *C*_*F*_-switch. However, this effect is only present for pulses in which there is a portion of the Pareto front for which the *U*-switch is superior, which only occur for relatively small amplitude pulses (Figure 3).

**Figure 5:**
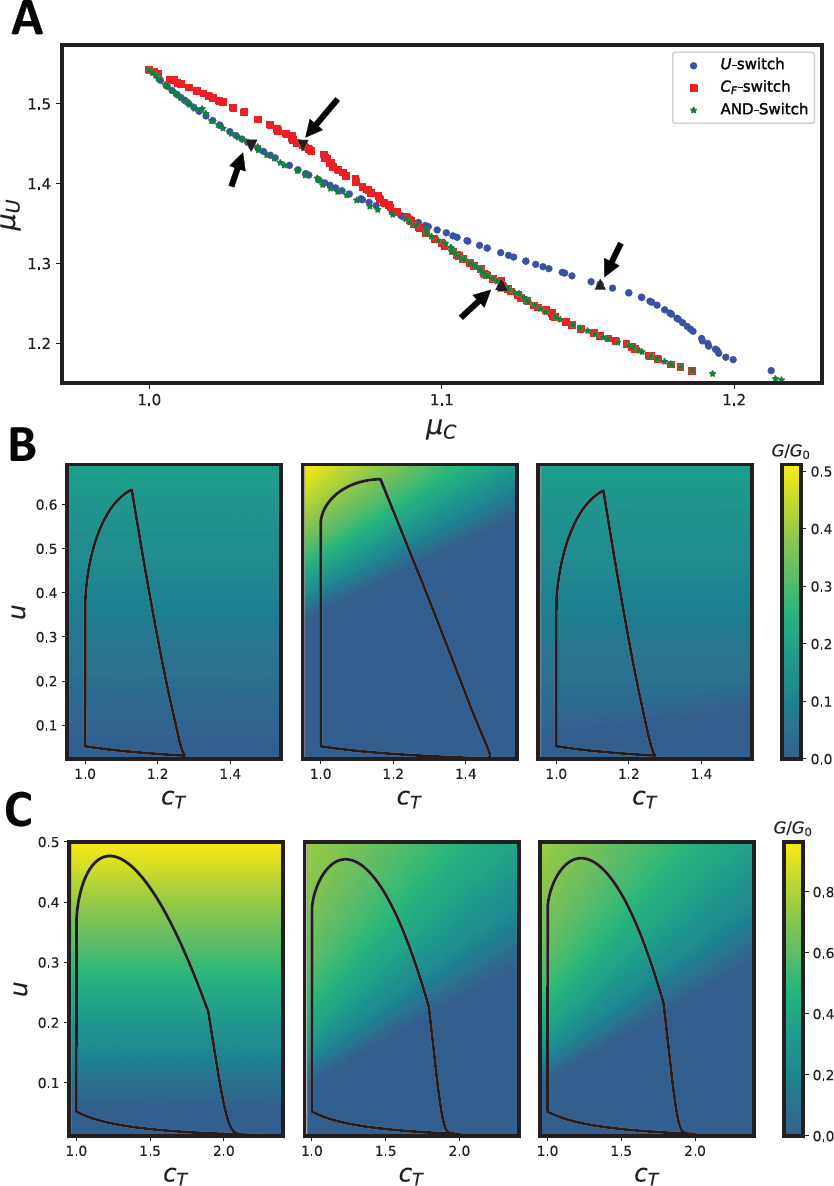
Comparison of Pareto fronts for AND-switch, *U*-switch and *C*_*F*_-switch. Panel A shows the Pareto fronts for the *U*-switch, *C*_*F*_-switch, and the AND-switch. The AND-switch coincides with the better of either the *U*-switch or the *C*_*F*_-switch in all cases, but does not out-perform the individual mechanisms. Panels B and C show the phase-plane trajectories and activation levels for sets of points on the Pareto fronts for which the AND-switch coincides with the *U*- switch (downward-pointing triangles) and the *C*_*F*_-switch (upward-pointing triangles). The left-most panels are for the *U*-switch, the center panels are for the *C*_*F*_-switch, and the right-most panel are for the AND-switch. The pulses parameters are: *D =*1.0, *τ*_*p*_ *=*0.5.

The preference for the AND-switch to coincide with either the *U*- or *C*_*F*_ -switch can be understood by considering the large-*m* limit in which both the *U*-switch and *C*_*F*_-switch become approximately binary. Then, the fully-active region of the UPR in phase space is where *u > u*_*min*_ and *C*_*F*_ < *C*_*F,max*_. Three scenarios are then possible for the control of activation (shown schematically in Figure 6), each of which depends on where the two activation thresholds intersect in the phase plane. First, if the intersection occurs at a value of *C*_*T*_ that is less than the steady state, then both the activation and deactivation thresholds will be determined by the *C*_*F*_ condition. In the second case, if the intersection occurs at a *C*_*T*_ that is greater than the steady-state chaperone concentration (since no change in total chaperone occurs until after the UPR is activated), but less than the maximal value reached during the stress event, the activation threshold will be determined by *u*_*min*_, and the deactivation threshold will be determined by *C*_*F*,*max*_. In the third case, if the intersection is located at a *C*_*T*_ value greater than the maximal level encountered by the system, then the control of both activation and deactivation depends only on *u*_*min*_. The first and thirds scenarios correspond to the *C*_*F*_-switch and *U*-switch, respectively, while the second scenario utilized both conditions of the AND-switch’s logic to separately control the thresholds for turning the UPR on and off. However, under the conditions shown in Figure 5, the optimal AND-switch always is of the type described in the first case. Yet, even though the activation threshold is set by the *C*_*F*_-switch, the slope of the response allows the activation surface (heat map in Figure 5B, right panel) to closely mimic that of the *U*-switch (left panel).

**Figure 6:**
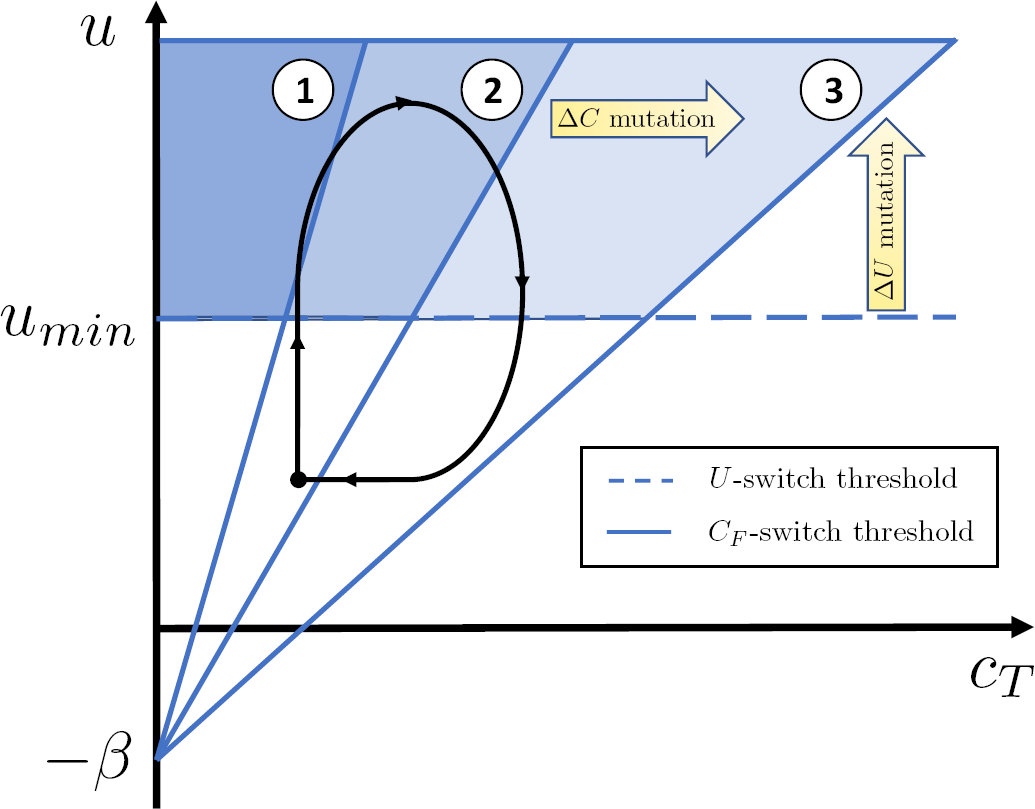
Schematic depiction of AND-switch phase-plane geometry. In the binary-switch limit, the intersection of the activation thresholds for the *U*-switch (dashed blue line) and the *C*_*F*_- switch (solid blue lines) with the phase-plane trajectory of the system (solid black curve) mark the points at which the UPR activates and deactivates. The shaded regions above the *u*_*min*_ line and to the left of the *C*_*F,max*_ lines demarcate the area of phase space for which the UPR is activated under the control of the AND-switch. Line 1 shows a case for which *C*_*F,max*_ dictates both the “on” and “off” transitions for the UPR. In the case of line 2, the “on” transition is determined by *u*_*min*_, and the “off” transition is controlled by *C*_*F,max*_. Lastly, line 3 shows a case for which the “on” and “off” transitions are set by *u*_*min*_. The arrows labeled *ΔC* mutant and *ΔU* mutant show the predicted change to the activation thresholds for a mutant in which the interaction between chaperone and sensor is disrupted (*ΔC*), and a mutant in which the unfolded protein-sensor affinity is decreased (*ΔU*). We emphasize that this picture is schematic and for any real system the dynamics (black curve) will necessarily depend on the threshold values for the UPR.

Taken as a whole, these results suggest that a sensing mechanism that incorporates both chaperone sequestration and direct unfolded protein binding does not improve efficiency of the feedback response beyond what can be achieved by either mechanism individually. Therefore, optimization of the trade-off between chaperone production and stress mitigation can rationalize the observation of a chaperone-based sensor, as discussed in the previous section, but not the combination of this mechanism with direct unfolded protein activation, suggesting that the combined mechanism observed in both yeast and higher eukaryotes serves another purpose.

## Discussion

Only with the appropriate design can a sensor network for the UPR efficiently regulate protein homeostasis in the cell. We developed a minimal model of the UPR that incorporates the stress of increased protein influx or increased protein misfolding within the ER, the role of folding chaperones in mitigating that stress, and the sensory network that controls the magnitude and timing of the transcriptional feedback. This model was then used in a genetic multi-objective optimization algorithm to determine the Pareto-optimal set of signal-transducing functions mapping the stress levels in the ER to response levels of chaperone transcription. Pareto optimization provides a useful structure for the analysis of regulatory mechanisms within the cell that must strike a balance between a few (or many) competing measures of fitness. In particular, it removes the subjectivity often required when choosing weights for different fitness functions to generate a single scalar fitness variable. Instead, calculation of the Pareto front allows for a clear understanding of the trade-offs constraining the fitness space of a phenotype.

In this work, we have applied this technique to the problem of maintaining protein homeostasis in the ER through the activation (and deactivation) of the UPR. Optimality was assessed with regards to two metrics: (i) the integrated level of unfolded protein over the course of the stressing event, and (ii) the excess production of chaperone over the course of the stress.

### What makes a good sensor network?

Analysis of the model provided insight into desirable traits for stress sensing networks. First, for chronic stress, in which the system has time to reach a new steady state (or limit cycle), the steepness of the activation function has two opposing effects: A greater slope suppresses oscillations of the feedback, but at the cost of looser overall control of the unfolded protein level across the operational range of the homeostat. Since experiments of UPR activation show non-oscillatory, dose-dependent responses to stress (Pincus *et al.*, 2010), it seems as though a more graded response that suppresses oscillations and allows for stable intermediate UPR levels has been selected.

Second, the model helped to identify a benefit of sensing stress through the concentration of free chaperone, as opposed to the concentration of unfolded proteins directly. A BiP-mediated activation was initially proposed for Ire1-based stress sensing due to the clear connection between BiP-Ire1 coimmunoprecipitation and suppression of the UPR with BiP overexpression (Dorner *et al.*, 1992; Bertolotti *et al.*, 2000; Okamura *et al.*, 2000). The analysis presented here provides a possible reason that a BiP-based mechanism would evolve. Sensing the level of free chaperone can both provide a sharper response during increasing stress, and increase the deactivation threshold as the system returns to basal functioning. Hence, it provides both faster “on’’ and “off’’ responses, allowing for a more efficient use of excess chaperone. The notion that one function of BiP is to accelerate the deactivation of the UPR is supported by experiments: yeast with an Ire1 mutant that does not bind BiP exhibits delayed deactivation of the UPR upon the removal of stress compared to wildtype (Pincus *et al.*, 2010). Interestingly, it has been demonstrated in yeast (Pincus *et al.*, 2010), and more recently in human (Karagöz *et al.*, 2017), that unfolded proteins within the ER lumen interact directly with Ire1, and are essential for full UPR activation. It has been proposed that unfolded proteins act as a ligand for activating the UPR, while BiP plays the role of a buffer (Oikawa *et al.*, 2007; Pincus *et al.*, 2010). Our analysis shows that the AND-switch logic requiring both inputs can decouple the threshold for activating the UPR from the point at which the UPR deactivates, but this does not enhance the efficiency of the response with regard to the trade-off between chaperone-production and stress-mitigation. Hence, the combined sensory mechanism is likely the result of another factor such as noise reduction or ligand selectivity.

### Effect of sensor mutations on signaling

Our model provides qualitative predictions regarding the impact mutations to Ire1 will have on the sensor activation and sensitivity to stress. An Ire1 mutant that does not bind BiP effectively increases *C*_*F,max*_ such that the *C*_*F*_-switch would always be active (Figure 6). In the case of the AND-switch, this means the activation and deactivation are both controlled by the unfolded protein threshold, *u*_*min*_. The early shutoff provided by the chaperone-sensing portion of the AND-switch is lost. In support of this, time-course experiments for UPR activation using an Ire1 mutant that does not bind BiP show an increased lag time between the removal of stress and deactivation of the UPR compared to wildtype Ire1 (Pincus *et al.*, 2010). Similar to more detailed UPR models (Pincus *et al.*, 2010), our minimal model predicts precisely this effect. Furthermore, our model shows that the origin of the BiP-mediated deactivation lies in the differing dependence on unfolded protein concentration of the *U*-switch and *C*_*F*_-switch. The fact that this has been experimentally observed in yeast lends credence to the idea that cells have evolved to take advantage of the enhanced efficiency of a BiP-mediated stress response.

Similarly, our model predicts that mutations causing a decrease in the affinity between Ire1 and unfolded proteins, effectively raising *u*_*min*_, could have one of two effects. If the mutation is severe enough to prevent the interaction altogether, the threshold for activation for the AND-switch would increase to an unsustainable level of stress, and the UPR would never activate. Alternatively, if the mutation raised *u*_*min*_ by a smaller amount, but enough so that the intersection of the *U*-threshold and *C*_*F*_-threshold moved from either case 1 or 2 to case 3 in Figure 6, then both activation and deactivation would be set by the unfolded protein concentration. In this case, we would again expect that the early shutoff provided by the BiP-Ire1 interaction would be lost. Mutational experiments disrupting the proposed unfolded-protein binding site on Ire1 have shown that the UPR is diminished for given drug-induced stress levels (Credle *et al.*, 2005), indicating that the affinity of Ire1-unfolded protein interaction can be modulated. It would be interesting to see whether certain mutations also remove the BiP-controlled early shutoff by increasing the threshold of unfolded protein required for activation to a level that renders the Ire-BiP interaction irrelevant.

### Other features can affect the fitness of stress signaling

While our model is far simpler than the many-faceted response of the true UPR, it captures essential aspects of the ER stress response, and allows for clear analysis of a subset of the evolutionary constraints governing the sensing mechanism of Ire1. Our model also provides novel insight into the role played by the stress-detecting network in optimizing the UPR. Of course, there are several questions that require further inquiry. For one, we have only considered isolated incidents of stress in this work. In reality, cells experience a dynamic continuum of different unfolded protein loads on the ER folding machinery. It remains to be established how switch design might be altered when the evolutionary driver is a distribution of protein fluxes into the ER. In this scenario, it is tempting to speculate that a graded response would be even more valuable, as it would allow for greater specificity in response over a range of stress levels. In a similar vein, the relatively small copy number of Ire1 molecules in the cell (≈ 200 (Ghaemmaghami *et al.*, 2003)) implies that there will be significant noise in any signal transmitted from the ER lumen to the nucleus. Further work will examine how noise and information transmission effect fitness of stress signaling.

Lastly, we note that the model presented here is not limited to describing the feedback of the UPR. It can readily be extended to any feedback control mechanism of enzymatic reactions in which either the substrate or enzyme act as a positive or negative regulator of enzyme production.

## Conclusions

In summary, we developed a minimal model of the UPR to understanding optimal design of the ER stress sensor network. The most important results of our analysis are: (i) A graded response will, in general, suppress oscillations in chronic stress conditions, at the expense of looser regulation of unfolded protein concentration in the ER, (ii) Sensors whose activity is downregulated by the amount of free chaperone can improve fitness by activating and deactivating at different levels of stress, (iii) Integrating signals from free-chaperone levels and unfolded proteins imbues the stress signaling network with an additional degree of freedom for tuning control of the UPR. However, with this extra degree of freedom does not enhance the fitness of the controller with regard to the trade-off considered here. By unraveling the advantages gained by indirect regulation of the ER stress sensor, our approach helps to understand homeostatic controllers in other biological contexts as well as guide sensor design synthetic biology.

